# Diverse temporal dynamics of repetition suppression revealed by intracranial recordings in human ventral temporal cortex

**DOI:** 10.1101/711523

**Authors:** Vinitha Rangarajan, Corentin Jacques, Robert T. Knight, Kevin S. Weiner, Kalanit Grill-Spector

**Author notes:** Co-Senior Authors (Kevin S. Weiner, Kalanit Grill-Spector).

## Abstract

Repeated stimulus presentations commonly produce decreased neural responses - a phenomenon known as repetition suppression (RS) or adaptation – in ventral temporal cortex (VTC) in humans and nonhuman primates. However, the temporal features of RS in human VTC are not well understood. To fill this gap in knowledge, we utilized the precise spatial localization and high temporal resolution of electrocorticography (ECoG) from 9 human subjects implanted with intracranial electrodes in VTC. Subjects viewed non-repeated and repeated images of faces with long-lagged intervals and many intervening stimuli between repeats. We report three main findings: (i) robust RS occurs in VTC for activity in high-frequency broadband (HFB), but not lower frequency bands, (ii) RS of the HFB signal is associated with lower peak magnitude, lower total responses, and earlier peak responses, and (iii) RS effects occur early within initial stages of stimulus processing and persist for the entire stimulus duration. We discuss these findings in the context of early and late components of visual perception, as well as theoretical models of repetition suppression.

## Introduction

Repeated exposures to sensory stimuli produce decreased neural responses, a phenomenon known as repetition suppression (RS), habituation, or adaptation. RS is common across sensory modalities in humans (Buckner RL et al. 1995; Grill-Spector K et al. 1999; Henson R et al. 2000; Alink A et al. 2018) and non-human primates (Gross CG et al. 1969; Gross CG et al. 1979; Miller EK et al. 1991; Lueschow A et al. 1994; McMahon DB and CR Olson 2007). RS is also considered a simple form of sensory learning (Thorpe W 1956; Groves PM and RF Thompson 1970; Miller EK *et al.* 1991) and a critical mechanism for perception (Grill-Spector K et al. 2006). Despite the ubiquity and proposed role of RS in perception, a surprising dearth of studies have examined RS using human intracranial measurements.

To our knowledge, the few studies that have examined RS in human VTC using intracranial electrophysiology implemented experimental designs in which the inter-stimulus intervals (ISIs) were short (largely between 275 and 500 milliseconds (Puce A et al. 1999; McDonald CR et al. 2010; Engell AD and G McCarthy 2014; Rodriguez Merzagora A et al. 2014; Miller KJ et al. 2015). These experiments were motivated by a) seminal findings showing that RS decreases with increasing ISIs, and is ablated once the ISI reached 20 seconds (Miller EK *et al.* 1991), and b) recent studies of RS in macaque inferotemporal cortex, which continue to use short ISIs to maximize RS (McMahon DB and CR Olson 2007; Verhoef BE et al. 2008; De Baene W and R Vogels 2010; Kaliukhovich DA and R Vogels 2011; Kuravi P and R Vogels 2017). Interestingly, event-related potential (ERP) and functional magnetic resonance imaging (fMRI) experiments have replicated and extended the characterization of repetition effects in the human brain (Schweinberger SR et al. 1995; Doniger GM et al. 2001; Schweinberger SR et al. 2002; Henson RN and MD Rugg 2003; Jacques C et al. 2007; Kuehl LK et al. 2013; Henson RN 2016). These studies illustrated that RS: a) occurs even with just a single stimulus repetition (Schweinberger SR *et al.* 2002; Sayres R and K Grill-Spector 2006), b) transpires across many intervening stimuli between image repetitions (Chao LL et al. 1999; Henson R *et al.* 2000; Sayres R and K Grill-Spector 2006; Weiner KS et al. 2010), and c) is larger for repetitions with no intervening stimuli and shorter time lags (Chao LL *et al.* 1999; Sayres R and K Grill-Spector 2006; Weiner KS *et al.* 2010; Kuehl LK *et al.* 2013).

Despite these findings across species and methodologies, the temporal characteristics of RS recorded directly from human VTC remain largely unknown, especially for long-lagged repetitions in which the repeated stimulus occurs after multiple other stimuli, typically after intervals longer than 10 seconds. Long-lagged repetitions are particularly interesting as they reflect an implicit neuronal memory trace and cannot be explained by refractory periods that may dampen the generation of action potentials (Grill-Spector K *et al.* 2006; Fabbrini F et al. 2019). As high-frequency broadband (HFB) signals are correlated with neuronal firing and local field potentials (Ray S and JH Maunsell 2011), recording HFB signals directly from the awake human brain offer a unique opportunity to determine the temporal characteristics of long-lagged RS.

To study the temporal characteristics of long-lagged RS, we conducted an electrocorticography (ECoG) experiment in 9 participants in which we measured electrical signals from VTC in response to novel and repeated images of faces. We examined repetition effects on visual responses to faces for two reasons. First, a large body of research documents both face-selective responses in VTC with ECoG (Allison T et al. 1999; McCarthy G et al. 1999; Parvizi J et al. 2012; Davidesco I et al. 2013; Rangarajan V et al. 2014; Jonas J et al. 2016) and robust RS for faces in VTC with fMRI (Grill-Spector K and R Malach 2001; Sayres R and K Grill-Spector 2006; Weiner KS *et al.* 2010; Engell AD and G McCarthy 2011). Second, all of our participants had electrodes located on the fusiform gyrus where face-selective regions reside (Kanwisher N et al. 1997; Weiner KS *et al.* 2010). In contrast, we did not have consistent coverage of other category-selective regions in VTC. We analyze both the magnitude and temporal characteristics of responses to repeated images and end the manuscript by discussing our findings in the context of theoretical models of RS (Grill-Spector K *et al.* 2006) as well as bottom-up and top-down components of visual perception.

## Methods

### Participants

9 subjects (3 female) were implanted with intracranial electrodes for neurosurgical evaluation for the treatment of refractory epilepsy. Electrode locations were chosen exclusively for clinical reasons by the subjects’ neurologist (JP) and neurosurgeons. The electrodes (AdTech) were implanted in the right and left hemispheres in 4 and 4 subjects, respectively, with 1 bilateral strip implantation. Electrodes had either 5 or 10 mm inter-electrode spacing (center to center) with an exposed recording surface diameter of 2.3 mm. All subjects gave informed written consent to participate in research studies at Stanford Medical Center as approved by the Stanford University Internal Review Board. All electrodes clinically implicated in the epileptogenic zone were excluded from analysis. In order to localize electrodes, high-resolution MRIs were acquired on a 3-Tesla GE scanner as previously described (Rangarajan V *et al.* 2014). Post-implantation computed tomography (CT) images were co-registered to the MRIs to visualize electrode locations in the subjects’ native head-space (Hermes D et al. 2010).

### Experimental paradigm (Figure 1)

Subjects participated in an experimental visual paradigm. Subjects viewed images of faces, limbs, cars, and houses that were presented foveally at a visual angle of approximately 10° by 10°. Each stimulus was presented for a duration of 1000 ms with a randomized inter-stimulus interval varying between 600-1400 ms. Some of the images were shown only once and some of the images were shown repeatedly (up to 6 times) during the experiment. On average, there were 8±1 intervening stimuli and 27.5±8.6 seconds between the 1^st^ and 2^nd^ presentations of the same image. The distributions of intervening stimuli and trial timing showed that 75% of trials had between 5-15 intervening stimuli and 10-30 seconds between repetitions. Images were equally likely to be repeated or nonrepeated throughout the course of the experiment. Images of all presented categories were included in both the nonrepeated and repeated conditions. The nonrepeated stimuli served two purposes (1) they were used as independent trials to assess the category selectivity of each electrode, and (2) they were used as intervening stimuli between repetitions of the same image.

**Figure 1:**
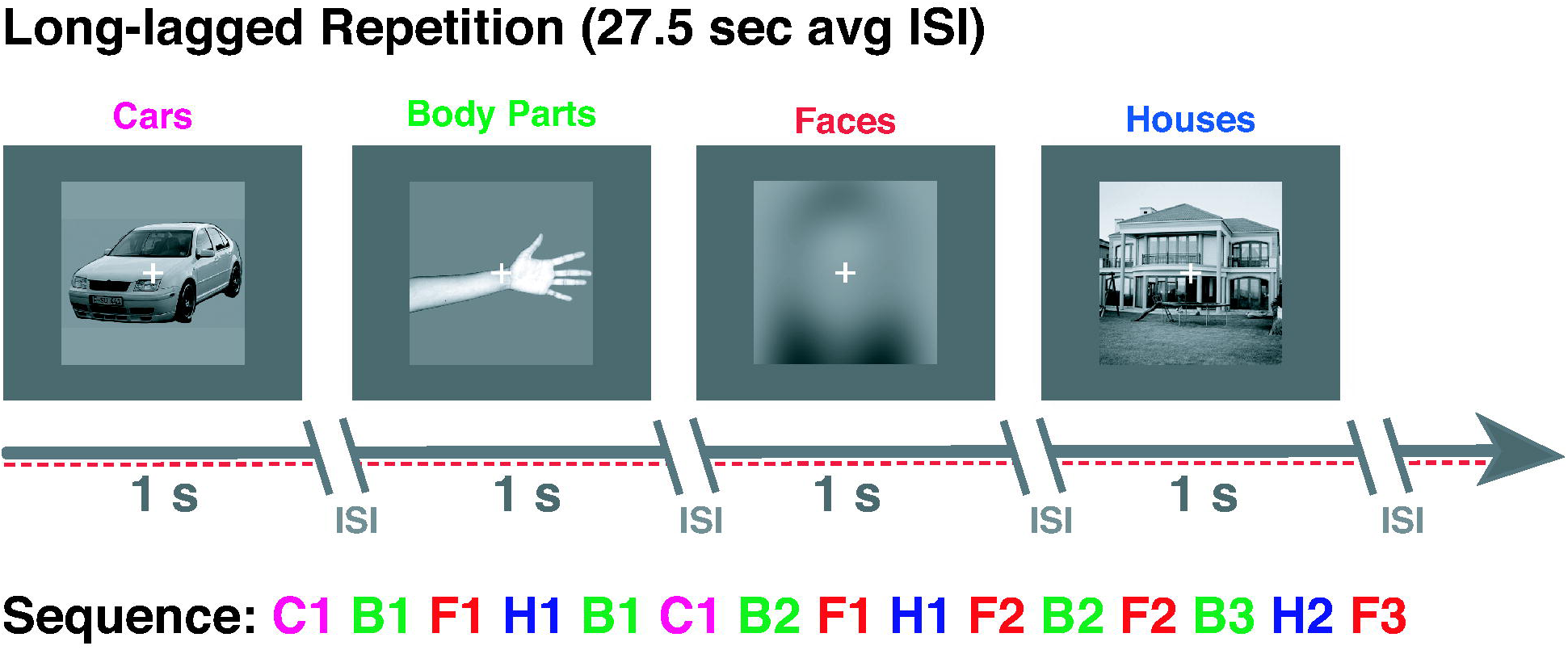
Design of repetition suppression (RS) long-lagged experiment. All 9 participants viewed images of cars, limbs, faces, and houses. Images were presented for 1 second each with a randomized inter-stimulus-interval (ISI) ranging between 600-1400 ms. The average time between repeats of the same image was 27.5 ± 8.59 seconds. *Task:* participants were instructed to fixate on a central cross and press a button on a keypad when it changed color from white to red. Example image sequences are shown for each experiment with colored letters indicating the stimulus category; *F:* faces; *L:* limbs; *C:* cars; *H:* houses. Numbers indicate the exemplar identity (e.g., F1 is face exemplar number 1).

### Task

Participants were instructed to fixate on a central fixation dot and to indicate when its color changed by pressing a button on an external keypad. Each subject participated in 5-10 runs of the experiment. Each run consisted of 96 stimuli which were divided into repeated (48 stimuli) and non-repeated (48 stimuli) images across the four stimulus categories. The order of repeated and non-repeated stimuli and order of categories was randomized for each subject with all 6 repetitions of an exemplar constrained to be contained within the same run. This resulted in an average of 16 repeated face exemplars per subject across runs (with each exemplar repeated 6 times each). One subject’s behavioral responses were not recorded because of a button-box malfunction (see behavioral performance in **Table 1**).

### Data Acquisition

Intracranial electrode activity was recorded on a Tucker Davis Technologies recording system sampled at 3052 Hz (subjects S1-7) and 1526 (S8-9). Signals were referenced online to a silent intracranial electrode and digitally bandpass filtered from 0.5 to 300 Hz. Pathological channels, as identified by the clinical team as showing seizure-like activity or falling within the seizure onset zone, and noisy (artifactual) channels were removed before data processing. Additionally, channels containing voltage greater than five times the mean variance in voltage were also removed. Finally, subsequent samples with greater than a 100uV jump, indicating an artifactual spike, were marked as bad temporal windows and excluded. The remaining channels (mean: 68 ± 26 electrodes/subject) were notch filtered at 60 Hz harmonics and then re-referenced to the common average to remove shared noise. The onsets of stimulus presentations were marked with a photodiode, time-locked to the neural data, and sampled at 24.4 kHz to maintain high temporal fidelity.

### Time-Frequency Data Processing

In order to be consistent with the literature in the field (Cole SR and B Voytek 2017; Kane N et al. 2017) and EEG/ECoG standards, the data were filtered for analysis in canonical frequency bands: theta (Θ: 4-8 Hz), alpha (α: 8-13 Hz), beta (β: 16-30 Hz), and high frequency broadband (HFB: 70-150 Hz). β and HFB data were obtained by bandpass filtering the original signal using 5 Hz non-overlapping frequency bins (e.g., 70-75, 75-80… 145-150 Hz). The time-varying power in each 5 Hz bin was estimated by applying a Hilbert transform (analytic amplitude squared). Each power timeseries was log transformed and the mean for the entire log power timeseries was subtracted. The transformed power timeseries were then averaged per canonical frequency band and downsampled to 100 Hz. This resulted in timeseries activity for Θ, α, β, and HFB activity corrected for the 1/frequency power decay in each frequency range. All analyses were done using custom MATLAB (Mathworks Inc) analysis scripts. All data were normalized (z-scored) with respect to the average of the 150 ms pre-stimulus baseline period per stimulus by subtracting the mean of the 150 ms before stimulus onset and dividing by the standard deviation of this time window.

### Localization of face-selective electrodes

In each subject, we identified face-selective electrodes using independent data from trials containing images that appeared only once during the long- lagged experiment, as in our prior studies (Parvizi J *et al.* 2012; Rangarajan V *et al.* 2014; Jacques C et al. 2016). In each electrode, we calculated the mean HFB power over a 0–900 ms window after stimulus onset averaged across all images from each category that were shown once. Then, we conducted two-sided t-tests to test if responses were significantly higher to images of faces compared to images of non-faces (p < 0.05). A false discovery rate procedure (FDR; Benjamini-Hochberg procedure; (Benjamini Y et al. 2001) for multiple-comparison correction was applied (α = 0.05) across electrodes within each subject. Importantly, the stimuli used for electrode-selectivity identification were excluded from all further analyses. Thus, all repetition suppression analyses were conducted with independent data.

To visualize the location of face-selective electrodes on a common MNI brain, the 3D whole brain anatomical images and electrode coordinates for each subject were converted to the MNI space and displayed over the mean cortical surface of the MNI brain (Hermes D *et al.* 2010) as in our prior studies (Rangarajan V *et al.* 2014). On the MNI brain, we displayed all recorded electrodes across subjects as well as the subset of face-selective electrodes, as illustrated in Figure 2. Given the variable nature of electrode coverage, we did not have many non-face category-selective electrodes (e.g. place-selective or limb-selective). In fact, no category, other than faces, had more than 10 selective electrodes across all 9 subjects (houses=7, limbs=10, cars=7).

**Figure 2:**
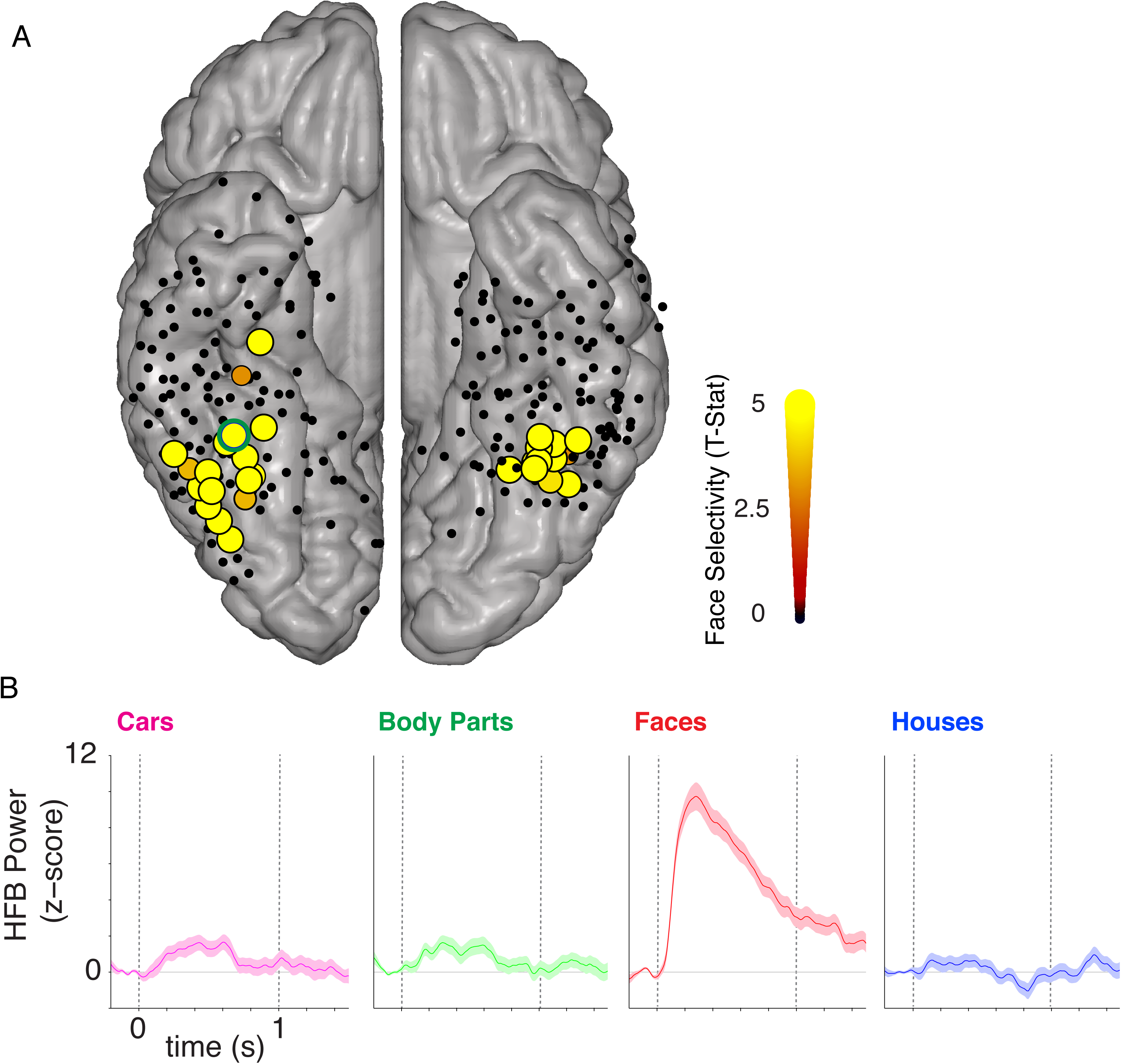
Face-selective electrodes in ventral temporal cortex. (A) All electrodes from the ventral temporal cortex (VTC) were projected into MNI space and illustrated on the Colin Brain cortical surface. The color and size of the electrode indicates the degree of face-selectivity, measured in the high frequency broadband (HFB, 70-150Hz) range. That is, larger size and brighter color indicates larger face selectivity (see colorbar). Face-selectivity was determined in each electrode by comparing the average magnitude of HFB response for faces versus non-faces during a time interval of 0 to 0.5 s. This selectivity was determined using independent trials during the long-lagged experiment which contained stimuli that appeared only once during the experiment. *Yellow electrodes* showed significantly higher HFB responses to faces than non-faces (t-value, FDR corrected p<0.05). *Black electrodes* did not show significant preference to faces. We identified a total of 48 face-selective electrodes in VTC, which were anatomically positioned on the fusiform gyrus (FG) and occipito-temporal sulcus (OTS) across the 9 subjects. On average, there were 6 ± 4 face-selective electrodes per subject. All subjects had at least 1 face-selective electrode. Some electrodes are not visible in the group average due to spatial overlap. (B) Average HFB responses across items of a category in an example electrode (green outline in A) located near the mid-fusiform sulcus (MFS). *Dashed vertical lines:* stimulus onset and offset, respectively.

### Analyses of neural responses to the 1^st^ and 2^nd^ presentation of images across frequency bands

Because different frequency bands are thought to represent varied features of electrocortical activity (Ray S and JH Maunsell 2011; Miller KJ et al. 2014), we examined responses to 1^st^ and 2^nd^ stimulus presentations separately for each of the canonical frequency bands: theta (Θ: 4-8 Hz), alpha (α: 8-13 Hz), beta (β: 16-30 Hz), and high frequency broadband (HFB: 70-150 Hz) in each electrode.

### Statistical analyses and justification of time windows

We performed a two-way repeated measures analysis of variance (ANOVA) using as factors: stimulus presentation number (1^st^ and 2^nd^ face presentations, this is a repeated factor, which resulted in 1 degree of freedom) and frequency bands (Θ, α, β, and HFB, which resulted in 3 degrees of freedom) across a time window from stimulus onset to the end of the trial (0-900 ms). The last 100 ms of the trial (900-1000) were omitted to avoid any edge effects resulting from the stimulus offset. Within-band statistical tests were done on the average response across all electrodes using t-tests comparing the total response for the 1^st^ and 2^nd^ face presentations over the entire (0-900 ms) window. To ensure any within-band effects were not due to the early evoked response alone, we next performed statistical analyses comparing responses across 1^st^ versus 2^nd^ presentations in two smaller time-windows: (i) 0-350 ms (reflecting early visually evoked responses) and (ii) 350-900 ms (reflecting the response after the evoked response)

### Determining significant electrode-level repetition suppression effects

Subsequent analyses focused on HFB activity because only HFB responses showed significant differences in responses between the 1^st^ and 2^nd^ presentations. Additionally, HFB activity is thought to reflect population spiking activity (Manning JR et al. 2009) and local post-synaptic activity (Logothetis NK et al. 2001; Miller KJ et al. 2010), which correlates with BOLD responses (Hermes D et al. 2012; Jacques C *et al.* 2016). To determine if RS was significant (p < 0.05), in each individual face-selective electrode we conducted a two-sided t-test of the response to the 1^st^ versus 2^nd^ face presentations in the 0-900 ms time window, which is reported in **Table 1**. A false discovery rate procedure (FDR: Benjamini-Hochberg procedure (Benjamini Y *et al.* 2001)) for multiple- comparison correction was applied (α = 0.05) across electrodes.

### Movie 1

To visualize the temporal dynamics of neural responses to stimulus repetition in VTC, we implemented a two-fold procedure. First, for each electrode, we computed a paired *t-*statistic comparing mean HFB responses for the 1^st^ versus 2^nd^ face presentation. These values were calculated in each 10 ms bin during an interval between 0-900 ms after stimulus onset. Second, we generated a movie of these values by visualizing the t-value of each electrode on the MNI surface across 10 ms time bins. This enabled visualizing the spatial and temporal effects of stimulus repetition across face-selective electrodes. For display purposes, the additive spatial distribution of t-values across electrodes were scaled to the same maximum and minimum (−2 (blue) and 2 (red), respectively), and corrected for the spatial distribution and overlap of included electrodes. In a single subject plot, electrode activities are plotted as a Gaussian around each electrode. With numerous subjects in MNI space, electrode coordinates overlap, which could falsely create patches of higher electrode density to appear to have more activity. To correct for this, we created a density map of the cortex, in which every electrode is given a value of 1. Anatomical areas with higher density of electrodes are then summed, and would have a higher value (e.g. 10). These numbers for electrode density are then normalized between 0 and 1 (so electrodes in the example high density region would be 0.1). This scaled density value is then multiplied by the activity of each electrode to correct for the spatial density of the channels, which controls for varied electrode coverage and prevents areas with higher spatial overlap of electrodes from appearing to be more face responsive. The video is slowed down 10 times such that 900 ms is displayed for 9 seconds. We tested if the average values for each 10 ms time window were significantly different using a t-test (1^st^ versus 2^nd^ presentation, p<0.05) to identify the rough onsets of RS or repetition enhancement effects at the group level.

### Metrics quantifying stimulus repetition

In all face-selective electrodes, we quantified the effect of stimulus repetition on HFB responses from single trials (e.g., responses to a single stimulus) using four metrics that capture the features of the HFB temporal response as well as its overall response amplitude. These effects were measured for face as well as house, limb, and car image repetitions.

1. Area under the response curve (AUC) is a measure of the total neural response. It was measured separately for each stimulus for the 1^st^ (non-repeated) and 2^nd^ (repeated) stimulus presentation. AUC was calculated for each trial using a trapezoidal numerical integration, which reflects a sum of all responses in the HFB band between a 0 to 900 ms time window in 10 ms steps per trial.
2. Peak Magnitude (PM) was calculated by finding the peak (maximum) HFB z-score value separately for the 1^st^ and 2^nd^ presentation of each stimulus and then measuring the average response across a 10 ms window surrounding the peak. The 10 ms time-window provides an estimate of the peak response that is less susceptible to noise-related fluctuations.
3. Response Onset Latency (ROL) of the neural signal was calculated by adapting the approach of Flinker A et al. (2010). We first identified the trials that showed a response to the stimuli. Responsive trials were defined as those trials in which responses significantly exceeded a null distribution of 1000 mean HFB epochs with random sampling. For these trials, we measured ROL during a 0–500 ms time window after stimulus presentation. The ROL was defined as the first time-point during the 0–500 ms time window in which the response passed the HFB z-score threshold (p<0.05 relative to baseline) for a duration of least 50 ms. This criterion was to ensure that we did not consider a momentary fluctuation as the response onset.
4. Peak Timing (PT) was calculated as the timing of the maximal (peak) response for a given trial. Peak response was defined as the maximal HFB z-score value. PM and PT were calculated separately for the 1^st^ and 2^nd^ presentation of each image. This trial-by-trial resolution enabled us to assess temporal variability between trials. All peaks >500 ms were excluded as noise for a conservative estimate of peak timing.

### Statistical testing

AUC, PM, ROL, and PT values were calculated at the single trial level, then averaged across trials per electrode. We examined the mean difference values (2^nd^ – 1^st^ presentation) per electrode as well as the total mean across electrodes for each of these metrics for 1^st^ and 2^nd^ presentations across all trials for each electrode. We then tested whether these values followed a normal distribution in order to determine which statistical tests were appropriate. Distributions that exhibited a skew of < ± 2 and a kurtosis < ± 7 were tested using parametric tests *(Bryne BM 2001; Hair JF 2009).* Here, we tested if the mean values across electrodes for each metric were significantly different across the 1^st^ versus 2^nd^ presentation using two-sided *t*-test to identify which metric showed a significant positive or negative shift (p<0.05). Those that were not normally distributed were tested with a nonparametric Wilcoxon rank sum test. All analyses were then repeated for the subsequent repetitions of each stimulus, i.e., 3^rd^, 4^th^, 5^th^, and 6^th^ presentations of faces.

### Assessment of repetition effects for non-preferred stimuli in face-selective electrodes

AUC, PM, ROL, and PT values were calculated at the single trial level for the non-preferred stimulus categories (houses, limbs, cars), then averaged across trials per electrode. We examined the mean difference values (2^nd^ – 1^st^ presentation) per electrode as well as the total mean across electrodes for each of these metrics for 1^st^ and 2^nd^ presentations across all trials for each electrode. Distributions that exhibited a skew of < ± 2 and a kurtosis < ± 7 were tested using parametric tests. Here, we tested if the mean values across electrodes for each metric were significantly different across the 1^st^ versus 2^nd^ presentation using two-sided t-test to identify which metric showed a significant positive or negative shift (p<0.05). Those that were not normally distributed were tested with a nonparametric Wilcoxon rank sum test. To correct for multiple comparisons, we used a false discovery rate procedure (FDR: Benjamini-Hochberg procedure (Benjamini Y *et al.* 2001)) with α = 0.05.

### Discriminability of distributed response to faces versus other categories over time and repetitions

To determine the representational structure and discriminability of faces from non-face categories, we measured distributed responses across face-selective electrodes. To remove between-electrode differences in overall amplitude that can be due irrelevant factors such as impedance, we normalized the time-course of each electrode. To do this, we concatenated the Training (non-repeated), Presentation 1, and Presentation 2 trials for all 4 categories (Face, House, Limb, Car) and then detected the top and bottom 10% of values per electrode. The mean of the top 10% and bottom 10% were set to the Maximum and Minimum, respectively. We then subtracted the minimum from each time point and divided it by the maximum minus minimum value ((value – min)/(max –min)) to normalize the electrode timecourse to a range between 0-1. This normalization was done separately for each electrode. One subject (S8) showed electrode timecourses that had outlier temporal responses across all conditions determined using the Thompson’s Tau outlier rejection (mean onset of electrode more than three standard deviations away from the mean onset across electrodes). These outliers generated a between-electrode difference that affects the pattern of distributed responses. To eliminate between-electrode effects in distributed analyses, we removed these outliers (6 face-selective electrodes of S8).

To determine the discriminability of patterns from each category, we then calculated the correlation between the distributed response across the 42 face-selective electrodes between the training set (non-repeated images), and each of the testing sets (Presentation 1 and Presentation 2) for each of the 4 stimulus categories (faces, houses, cars, and limbs). Discriminability is defined as significantly higher within-category than between-category correlations. Correlations between distributed responses were calculated in 10 ms time windows from 150 ms before stimulus onset till 900 ms after stimulus onset. For each pairwise correlation, we calculated the bootstrapped (50 iterations) correlation across 75% of electrodes (n=36 electrodes) that were randomly selected with replacement, to allow for an error estimate on the correlation values. We reasoned that face information would be associated with higher within-category distributed responses (face-face) than between-category distributed responses (limb-face/car-face/house-face). To evaluate if face-discriminability arises in different times for Presentation 1 versus Presentation 2, we computed the within-category (face-face) correlation minus the average of between-category correlations (face/non-face) for the first and second presentations in each 10 ms time window with bootstrapping (50 iterations, 75% electrodes per bootstrap) and tested if discriminability varied across repetitions and at which time bins.

## Results

### Stimulus repetition reduces neural responses (repetition suppression, RS) in high-frequency broadband, but not other frequency bands

We first examined the effect of long-lagged image repetitions on ECoG responses by comparing ECoG signals for the 1^st^ and 2^nd^ presentation of faces in each of the canonical Θ, α, β, and HFB frequency bands. Figure 3 shows the mean response across all 48 face-selective electrodes, which were localized with independent data (Figure 2) for each of these frequency bands during the 1^st^ (red) and 2^nd^ (blue) presentation of face images (S1= 9, S2= 11, S3= 3, S4= 10, S5= 2, S6= 2, S7= 1, S8= 8, S9= 2 face-selective electrodes). Notably, we observed reduced responses for the 2^nd^ compared to the 1^st^ presentation (or repetition suppression, RS) in the HFB, but not other frequency bands.

**Figure 3:**
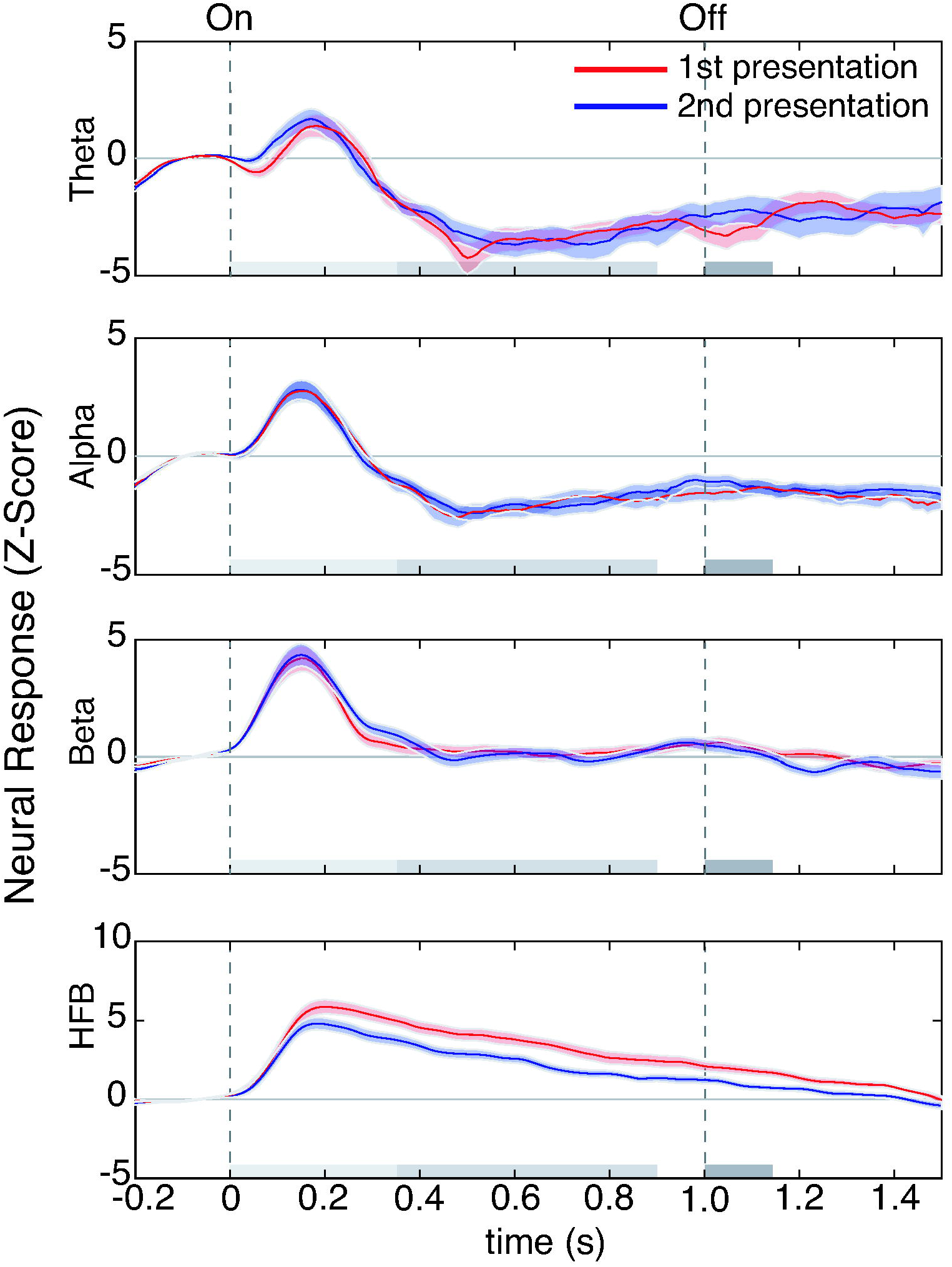
Repetition suppression (RS) in human VTC occurs in HFB, but not lower frequency bands. Average responses to 1^st^ (red) versus 2^nd^ (blue) presentations of faces across all face-selective electrodes. Each row shows neural responses in a different frequency band. *From top to bottom*, frequencies are: theta (Θ: 4-8 Hz), alpha (α: 8-13 Hz), beta (β: 16-30 Hz), and high frequency broadband (HFB: 70-150 Hz). *Shaded region:* standard error of the mean across 48 face-selective electrodes. *Dashed vertical lines*: stimulus onset and offset, respectively. *Gray horizonal bars* illustrate the following time-windows of interest: (i) *light gray:* 0-350 ms (evoked response), (ii) *medium gray:* 350-900 ms (later response), (iii) *light and medium gray*: 0- 900 ms (stimulus duration), and iv) *dark gray:* 1000-1150 ms (post-trial activity).

We assessed the significance of our observations using a two-way repeated-measures analysis of variance (ANOVA) on the total HFB response integrated over a 0-900 ms time window after the stimulus onset using presentation number (1^st^/2^nd^) as a repeated factor and frequency band (Θ, α, β, and HFB) as a factor. Across all subjects (S1-9), we found a significant main effect of frequency band [F(3, 188)=95.537, p=1.34×10^−37^], a significant main effect of repetition [F(1, 188)=11.712, p=7.6×10^−4^], and a significant interaction [F(3, 188)= 89.856, p<4.1x^−36^].

Given the significant main effect of frequency band and repetition, as well as the significant interaction, we conducted post-hoc analyses to identify which specific frequency bands and time windows were driving the effect. We first conducted this analysis across the whole time window used in the analysis above: 0-900 ms (stimulus duration). Subsequently, we wanted to ensure that any significant results were not simply due to the evoked response and therefore, used two smaller time windows: a time window from 0-350 ms (consisting of the evoked response), as well as a time window between 350-900 ms (the remainder of the stimulus duration after the evoked response).

Examination of the time course of neural responses revealed significant RS to the 2^nd^ versus 1^st^ presentation in the HFB response (Figure 3). RS was significant for HFB activity from the entire 0-900 ms window (t-test, t(47)=4.16, p=1.35×10^−4^, df=47: n-1 electrodes, FDR corrected). Significant RS in HFB was already evident in the early (0-350 ms) portion of the trial (t-test, t(47)=3.82, p=3.9×10^−4^, FDR corrected). This 0-350 ms temporal window is associated with the synchronized evoked visual response that is prominent in low frequencies. Notably, significant RS in the HFB persisted beyond the evoked response in the 350-900 ms time window (t-test, 350-900 ms; t(47)=4.12, p=1.5×10^−4^, FDR corrected). Interestingly, even up to 150 ms after the stimulus was off, RS persisted: there was a significant difference between the response to the 2^nd^ versus 1^st^ presentation in the 1000-1150 ms time window (t-test, t(47)=3.72, p=3.2×10^−4^, FDR corrected).

In comparison to neural responses in the HFB, responses in Θ, α, and β frequency bands were more transient. Specifically, responses in these frequency bands increased from baseline during a 0-350 ms time-window after stimulus onset, and then returned to baseline. Interestingly, there was no significant RS in Θ (t-test, t(47)=0.19, p=0.85, FDR corrected), α (t-test, t(47)=0.08, p=0.93, FDR corrected) or β (t-test, t(47)=0.11, p=0.91, FDR corrected) frequency bands across the entire stimulus duration (0-900 ms) or in the early evoked response period (t-test: Θ: t(47)=0.61, p=0.55; α: t(47)=-0.20, p=0.84; β: t(47)=0.93, p=0.36). Altogether, these results indicate that RS largely occurs in HFB, but not lower frequency bands and that RS persists throughout the entire stimulus duration.

### Visualizing the spatiotemporal dynamics of RS in human VTC

We visualized the spatiotemporal dynamics of RS in human VTC. To do so, we generated **Movie 1** that illustrates the average difference in HFB response (t-values) between the 1^st^ and 2^nd^ presentation of each face image at 10 ms time bins for each of the 48 face-selective electrodes. Consistent with our subsequent analyses and statistical quantification, **Movie 1** shows that the most pervasive effect of stimulus repetition is reduced HFB responses to the 2^nd^ compared to the 1^st^ presentation. Spatially, this reduction (illustrated in the warm colors in **Movie 1**) occurs primarily in the lateral fusiform gyrus (FG). Interestingly, RS is evident early in the response, and appears within the first 100-200 ms after stimulus onset in several electrodes.

The first significant 10 ms window across all electrodes showing RS was on average 153.75 ms (90% CI: 23:410). RS a) persists throughout the duration of the stimulus, b) increases over time, and c) spreads both more posteriorly and anteriorly, particularly in the right hemisphere. Nevertheless, since there are only three left hemisphere electrodes with time bins (epochs) showing significant RS, we were not able to assess hemispheric lateralization effects. We also note that three electrodes in the right hemisphere showed a weak enhanced response to the 2^nd^ versus 1^st^ presentation. Of these, two electrodes were located in the right inferior temporal gyrus (ITG), and one was located in the anterior and medial aspect of the right FG. In these three electrodes, the first 10 ms time window showing significant repetition enhancement appeared later in the trial, on average 376.25 ms (90% CI: 26:690) after stimulus onset. The spatial and temporal dynamics of RS in HFB are further quantified in the following sections.

### Reduced total and peak response amplitudes in HFB for repetitions in single electrodes

The prior quantitative analyses of RS examined neural responses integrated across trials and across electrodes, which may obscure finer temporal characteristics of RS visible in **Movie 1**. Thus, we next examined the spatial and temporal dynamics of HFB responses in each electrode. As RS effects in the HFB persisted throughout the trial, all subsequent analyses were done in the 0-900 ms time window. To characterize responses over time, we calculated and compared four metrics of the HFB response for 1^st^ and 2^nd^ face presentations (Methods, Figure 4A): (1) total response (area under the response curve, AUC), (2) peak magnitude (PM), (3) response onset latency (ROL), and (4) peak timing (PT). Each metric was calculated at the single trial level and then averaged across all trials for each electrode. Statistics were performed across electrodes.

**Figure 4:**
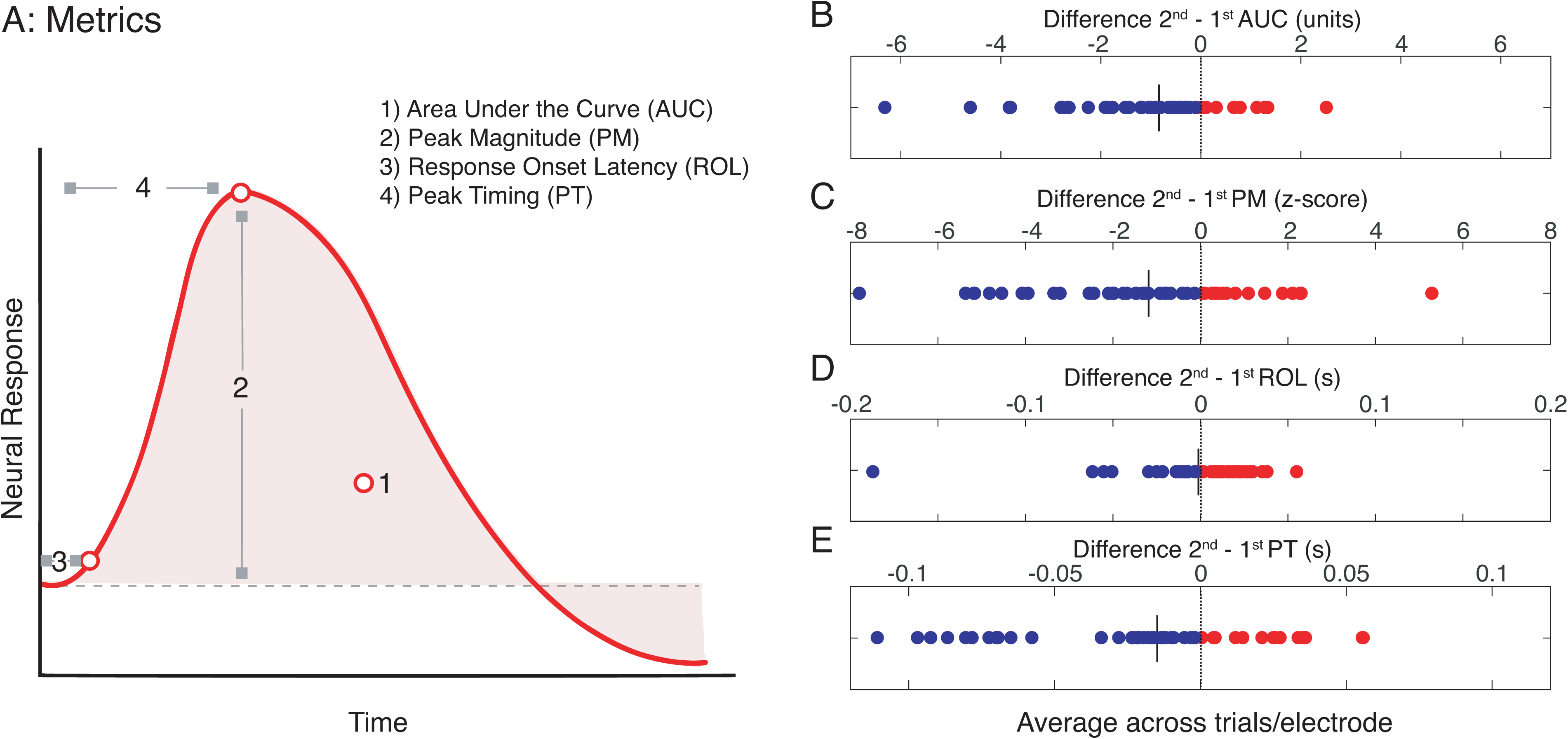
Quantifying the effect of repetition on the magnitude and timing of HFB responses. (A) Schematic illustration of four metrics of the HFB signal that were calculated for each of the 48 face-selective electrodes: (1) Total HFB response, which is the area under the response curve (AUC) from 0-900 ms (light red), (2) peak magnitude (PM), (3) response onset latency (ROL), and (4) peak timing (PT). (B) Difference in total response, area under curve: AUC (2^nd^ presentation) – AUC (1^st^ presentation) for faces. (C) Difference in peak magnitude: PM (2^nd^ presentation) – PM (1^st^ presentation) averaged over a 10 ms window surrounding the peak. (D) Difference in response onset latency: ROL (2^nd^ presentation) – ROL (1^st^ presentation). (E) Difference in peak timing: PM (2^nd^ presentation) – PM (1^st^ presentation). In B-E, each point is an electrode. *Blue:* negative values indicate decrements. *Red:* positive values indicate increments. *Vertical line:* mean value across electrodes.

Examination of the effect of repetition on (a) the total HFB response (AUC) and (b) the peak magnitude (PM), revealed two main findings. First, the total HFB response was reduced for 2^nd^ versus 1^st^ presentation of faces at the fine-spatial scale of a single electrode (Figure 4B). The mean AUC for the 1^st^ presentation was 3.4 ± 2.1 versus 2.5 ± 1.6 for the 2^nd^ presentation. 77.1% (37/48) of electrodes across all (9/9) subjects showed lower AUC values for the 2^nd^ versus 1^st^ presentation (**Table 1**), leading to a negative difference in the AUC between presentations (blue dots in Figure 4B) at the group level. Statistical comparisons of the 1^st^ versus 2^nd^ presentations showed a significant reduction in the total AUC of HFB responses (Figure 4B, paired t-test t(47)= 4.16, p=1.34×10^−4^, FDR corrected). Additionally, 22.9% (11/48 electrodes from 6/9 subjects) of electrodes showed a change in the AUC that is either equal to zero or enhanced (2^nd^ > 1^st^). The specific electrodes showing a significant reduction in total response (paired t-test, FDR corrected) are indicated in **Table 1**. In contrast, only 3 electrodes showed a significant repetition enhancement in the HFB reflected by a substantial increase in the AUC (paired t-test p<0.05, FDR corrected, Supplemental Figure 1). Further, repetition enhancement occurred only in smaller intervals during the trial, but not over the complete trial (**Movie 1**).

Second, we examined the effect of repetition on the peak magnitude (PM) of responses in each electrode. PM metrics are reported in units of z-score relative to the baseline (mean pre-stimulus response) to control for between-electrode differences in overall response magnitude (see Methods). This analysis revealed that PM of the response was smaller for the 2^nd^ compared to the 1^st^ presentation of a face. Indeed, the mean PM for the 1^st^ presentation was 7.5 ± 3.0 standard deviations away from the baseline, while the 2^nd^ presentation was only 6.3 ± 2.6 standard deviations away from the baseline. Similar to the analysis of the AUC, the majority of electrodes (64.6%, 31/48 electrodes from 8/9 subjects) showed a negative PM difference between the 2^nd^ versus 1^st^ presentation (Figure 4C, paired t(47)=3.39, p=0.001, FDR corrected). PM reduction between 2^nd^ versus 1^st^ presentations was significantly correlated with AUC decrease between 2^nd^ – 1^st^ presentations of faces (Pearson *r*=0.87, p=4.6×10^−16^). These analyses suggest that both AUC and PM reduce with repetition and these magnitudes are significantly correlated.

One may intuit that changes in task performance may contribute to reductions in PM or AUC for the second presentation. To remind the reader, subjects were instructed to indicate a change in the color of the central fixation dot. We note, however, that task performance cannot explain reductions in PM or AUC, as the changes in fixation did not occur on every trial, and were orthogonal to both the repetition and content of our stimuli (**Table 1**). Together, these analyses indicate that both the total HFB response and peak HFB response are lower for the 2^nd^ versus 1^st^ presentation of a face, illustrating significant RS in face-selective electrodes for long- lagged stimulus repetitions.

### Peak timing is faster for the 2^nd^ compared to the 1^st^ image presentation in a majority of electrodes

Next, we quantified the temporal profile of neural responses to face repetitions using two metrics: (1) the response onset latency (ROL) of HFB responses, and (2) peak timing (PT) of HFB responses. The mean response onsets to the 1^st^ and 2^nd^ presentations were 101.8 ± 38 ms and 100.2 ± 31.4 ms, respectively, which were not significantly different. Examination of the differences in ROL for the 2^nd^ versus 1^st^ presentations of faces showed a delayed onset in 27/48 electrodes (56.3%, 8/9 subjects) for 2^nd^ versus 1^st^ presentations of faces, while the remaining electrodes (43.7%, 21/48 electrodes from 8/9 subjects) showed the opposite effect (Figure 4D). Consequently, across electrodes, there was no significant difference in the ROL (Wilcoxon rank sum test, z= −0.726, p=0.468) between the 1^st^ and 2^nd^ presentations.

It is interesting that while there were no differences in ROL, PT was earlier for the 2^nd^ compared to the 1^st^ presentation of a face in the majority of face-selective electrodes (64.6%, 31/48 electrodes, 8/9 subjects). Indeed, the mean PT for the 1^st^ presentation of a face was 276 ± 39 ms and for the 2^nd^ presentation was 259 ± 51 ms. Statistical comparisons of the 1^st^ versus 2^nd^ presentations showed a significant earlier PT (paired t-test: t(47)=2.83, p=0.006, FDR corrected), which was on average 17 ± 4 ms faster for the 2^nd^ than 1^st^ presentation. Additionally, a minority of electrodes (n=5) showed the converse pattern and a significantly delayed PT (p<0.05) during 2^nd^ compared to 1^st^ stimulus presentations. Notably, PT did not correlate with either AUC (Pearson r=0.18, p=0.23) or PM (Pearson r=0.065, p=0.66). This seems to indicate that while the timing of the peak is affected by repetition, it does not correlate with the magnitude-based metrics of RS.

To confirm that our findings were not due to outliers, we implemented a Thompson’s Tau outlier rejection which removes samples more than three standard deviations away from the mean. The outliers removed were not all from the same electrode and the group comparisons of AUC, PT, and PM between 1^st^ and 2^nd^ presentation remained significant. Taken together, analyses of the temporal dynamics illustrate that HFB responses are overall lower, have a smaller peak magnitude, as well as an earlier time to peak for the 2^nd^ compared to the 1^st^ presentation of faces.

### RS and temporal dynamics are different for non-preferred categories compared to faces in face-selective electrodes

An interesting question is whether RS also occurs for non-preferred stimuli in face-selective electrodes. Prior fMRI data show that the magnitude of RS (difference in signal amplitude) is larger for the preferred category than the non-preferred category, consistent with a scaling mechanism of RS (Weiner KS *et al.* 2010). Thus, we next examined the responses to the 1^st^ versus 2^nd^ presentation of houses, limbs, and cars in the same face-selective electrodes (electrodes shown in Figure 2). We tested if changes to the response profile for 2^nd^ versus 1^st^ presentations of non-preferred categories were similar to the RS effects (lower total response (AUC), smaller peak magnitude (PM), and earlier peak timing (PT) observed for 2^nd^ versus 1^st^ face presentations. We emphasize that we only examined RS effects to other categories within face-selective electrodes as opposed to also examining RS to images of these categories in other electrodes that may be selective to these categories (e.g., house repetitions in house-selective electrodes) due to limited electrode coverage in VTC (Methods).

The largest effect of RS was on the total response change for 2^nd^ versus 1^st^ presentations of faces. When comparing the total response (AUC) between the 1^st^ versus 2^nd^ presentation of houses, limbs, and cars, only repetitions of limbs caused a significant reduction (AUC paired t-test: t(47)=2.23, p=0.03, FDR corrected), while cars (Wilcoxon rank sum: z=-0.3, p=0.76) and houses (t-test: t(47)=1.78, p=0.08, FDR corrected) did not (Figure 5A). This lack of RS may be due to the limited initial HFB response to the 1^st^ presentation of cars and houses in face-selective electrodes (see example responses in Figure 2). However, some electrodes show an enhancement effect for houses where the 2^nd^ presentation elicits a higher HFB response than the 1st.

**Figure 5:**
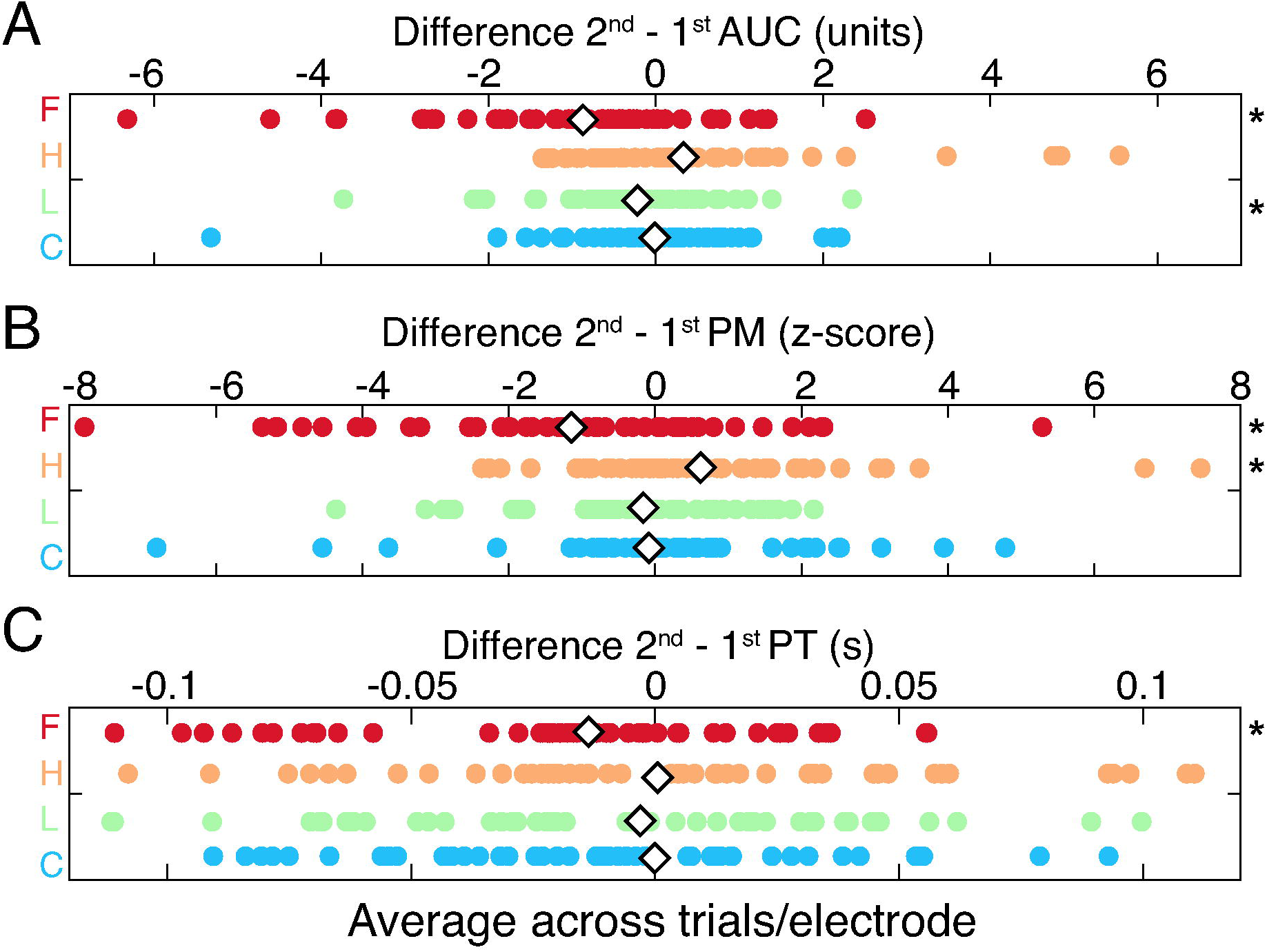
Repetition effects for non-preferred categories in face-selective electrodes. The mean difference between 2^nd^ – 1^st^ presentations for (A) Area under curve, AUC, (B) Peak magnitude, PM, and (C) peak timing, PT. Each dot indicates a single electrode value. *Red*: F, faces; *orange:* H, houses; *green:* L, limbs; *blue:* C, cars. Asterisk: significant difference (p< 0.05, t-test) between 1^st^ versus 2^nd^ presentation of the face, FDR corrected across electrodes. *White diamond*: mean value for each presentation number.

Similarly, when comparing the change in PM to the 1^st^ versus 2^nd^ presentation of houses, limbs, and cars, there was no significant reduction in PM between 2^nd^ versus 1^st^ stimulus presentations in face-selective electrodes (Figure 5B, limbs paired t-test: t(47)=1.39, p=0.17; cars Wilcoxon rank sum test: z= −0.64, p=0.76). In fact, the PM was actually enhanced in house repetitions (paired t-test: t(47)=2.18, p=0.03, FDR corrected) indicated by the rightward shift of the house difference values in Figure 5B. This suggests that in face-selective electrodes, a reduction occurs for the preferred stimulus (faces), while there is largely no-change or enhancement in response to non-preferred stimulus categories.

Finally, we compared the changes in timing parameters (ROL and PT) to the 1^st^ versus 2^nd^ presentation of houses, limbs, and cars in face-selective electrodes. As no ROL change was found for face repetitions, no change was expected for ROL for repetitions of images from non-preferred categories as well. Houses (paired t-test: t(47)=1.64, p=0.1075, FDR corrected) and limbs (paired t-test: t(47)=2.04, p=0.05, FDR corrected) showed no change for the ROL, but cars showed a slightly earlier response onset for the 2^nd^ presentation versus the 1^st^ (paired t-test: t(47)=2.33, p=0.02, FDR corrected). As PT was earlier for the 2^nd^ presentation of faces in face-selective electrodes, we predicted an earlier PT for repetitions of non-preferred categories in these electrodes (Figure 4D). However, contrary to this prediction, there was no significant change to the PT for repetitions of houses (paired t-test: t(47)=1.16, p=0.25, FDR corrected), limbs (paired t-test: t(47)=1.92, p=0.06, FDR corrected), and cars (paired t-test: t(47)=0.786, p=0.436, FDR corrected) in face-selective electrodes (Figure 5C). This may indicate that temporal changes in the peak timing only occur robustly for repetitions of stimuli from the preferred category.

Overall, repetitions of images from non-preferred categories produced minimal to no RS. The only significant effects were a small reduction in the total response to limbs, some enhancement in the peak magnitude of houses, and a slightly earlier response onset for cars. The lack of consistent response changes to repetitions of non-preferred stimuli indicates that RS effects in face-selective electrodes are prominent for the preferred stimulus category (faces), but are limited/lacking for non-preferred categories.

### Discriminability of distributed response to faces versus other categories over time and repetitions

As we found earlier PT for repeated faces, but not other categories, in face-selective electrodes, we further investigated whether the discriminability of faces from non-face categories varied across repetitions. Thus, we computed face-discriminability of distributed responses across face-selective electrodes over. To estimate face-discriminability in an unbiased way, we used the non-repeated images from each category as a training set, and the distributed responses from the 1^st^ (Presentation 1) and 2^nd^ (Presentation 2) of each category as testing sets. We measured the correlation between the distributed pattern of response across face-selective electrodes for non-repeated images and the distributed patterns of response for Presentation 1 and Presentation 2 of repeated images. Distributed responses were calculated separately for each condition and category in 10 ms time bins using a bootstrapping approach (Methods, Figure 6A,B, Supplemental Figure 2). Then, we calculated the differences between the within versus between category correlations and examined their timing profiles.

**Figure 6:**
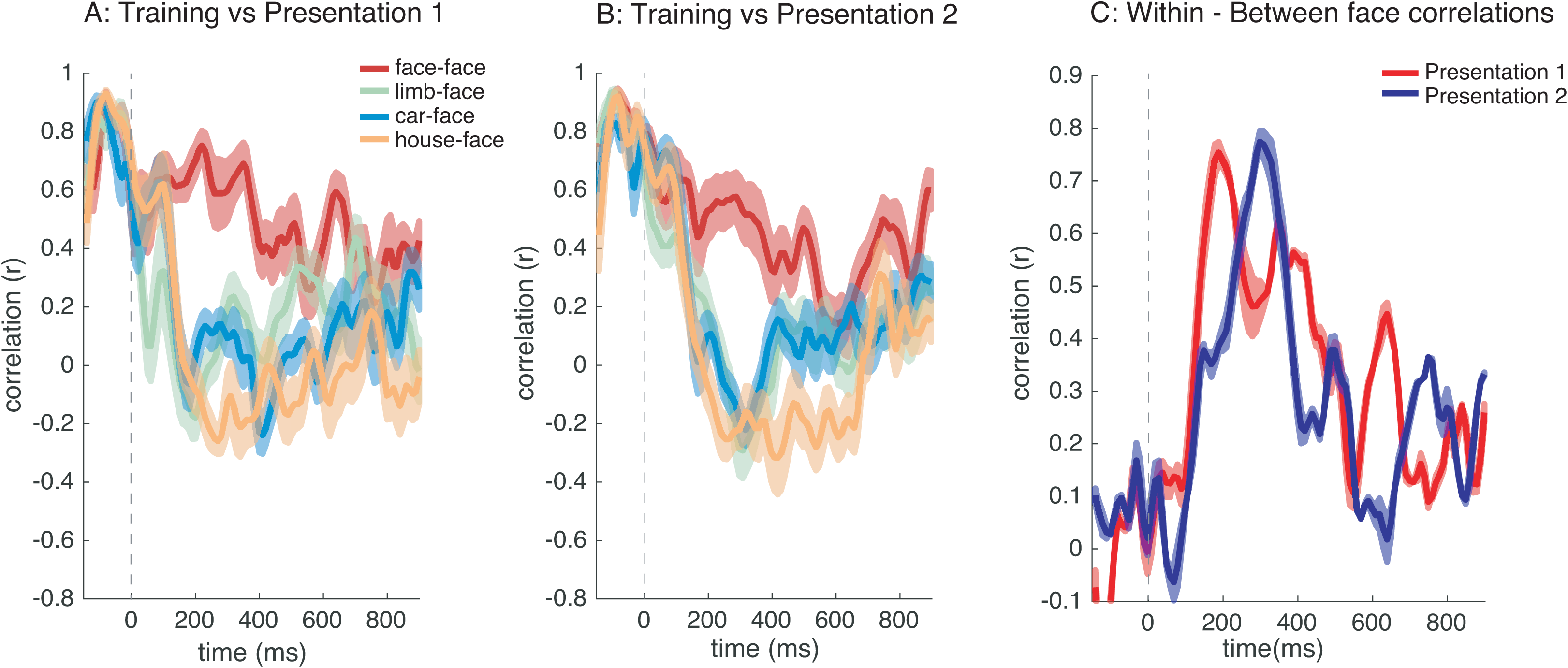
Analysis of distributed responses to the four categories (face, limb, car, house) over time and repetitions. The correlation between the distributed responses of training images (non- repeated faces) and each of the testing sets across 42 electrodes: (A) 1^st^ and (B) 2^nd^ face and non- preferred categories. Color indicates the categories that are correlated: *red:* face-face; *orange:* houses-face; *green:* limbs-face; *blue:* cars-face correlations, respectively). (C) Within-face correlation minus between mean face and non-face correlations for 1^st^ and 2^nd^ presentations are plotted over time; dashed vertical lines: stimulus onset. *Error bars: indicate standard error across bootstraps.*

We found that for both Presentation 1 (Figure 6A) and Presentation 2 (Figure 6B) distributed responses across face-selective electrodes had the highest positive correlation to distributed responses to non-repeated faces from the training set compared to all other non-preferred categories (houses, cars, and bodies). This difference was most prominent starting at ˜200ms after stimulus onset. To evaluate face-discriminability, we measured the difference between the within-category correlation (face-face) and the mean of the between-category correlation (average of face-limb, face-car, and face-house) for each of the first and second presentations in each 10 ms time window. A significant positive difference indicates face- discriminability as it shows that within-category correlations are higher than between-category correlations.

Examining the temporal progression of discriminability of distributed response to faces (Figure 6C), shows that face discriminability is low within the first 100 ms after stimulus onset for both Presentation 1 and Presentation 2; face-discriminability then steadily increases till about ~200-400 ms after stimulus onset, where at that point face discriminability is positive and high (peak discriminability ~0.7). This data shows that there is face information in distributed responses across face-selective electrodes for both 1^st^ and 2^nd^ presentation. While the rise in discriminability is similar for the 1^st^ and 2^nd^ presentations, the peak discriminability is earlier for the 1^st^ than 2^nd^ presentation (Figure 6C), despite the fact that the PT was earlier for the latter than the former. The later peak in face discriminability for the 2^nd^ presentation is likely due to later lower between-category correlations to the 2^nd^ presentation compared to the 1^st^ presentation (Figure 6A/B). A winner-take-all classification analysis showed ceiling face classification for both 1^st^ and 2^nd^ presentations, and did not provide sufficient sensitivity to determine differences in classification timing across repetitions. Overall, discriminability of distributed responses to faces started to rise at a similar time for both the 1^st^ and 2^nd^ presentations, approximately 100ms after stimulus onset.

### RS magnitude and timing effects persist for additional repetitions of faces

Within our experimental design, each of the repeated images was presented 6 times, which allowed us to further investigate how additional repetitions of faces affect the HFB response. In the following analyses, we focus on responses to additional repetitions of faces because face-selective electrodes did not show reliable and robust RS effects for non-preferred categories. The average time course for each of the 6 face repetitions is shown in Supplemental Figure 3. Based on fMRI measurements of VTC responses to stimulus repetitions in which the level of neural responses typically asymptote for more than 4 repetitions (Sayres R and K Grill-Spector 2006), we predicted that HFB metrics related to response amplitude (AUC and PM) will decrease with additional repetitions of a face, but may asymptote after the 4^th^ repetition. We highlight, however, that neither fMRI data, nor theoretical models of RS, make strong predictions regarding the effect of additional repetitions on temporal features of the response such as the ROL and PT.

We first assessed the significance of our observations using a repeated-measures analysis of variance (ANOVA) on the total HFB response integrated over a 0-900 ms time window after the stimulus onset using presentation number (1^st^ through 6^th^) as a repeated factor. Across all subjects (S1-9), we found a significant main effect of repetition [F(5, 47)=7.19, p=2.78×10^−6^] because presentations 2-6 of a face generated lowered responses relative to its 1^st^ presentation (see Figure 7A for AUC). Given this significant main effect of RS, we calculated AUC, PM, ROL, and PT for the 3^rd^, 4^th^, 5^th^, and 6^th^ presentation of faces for each electrode.

**Figure 7:**
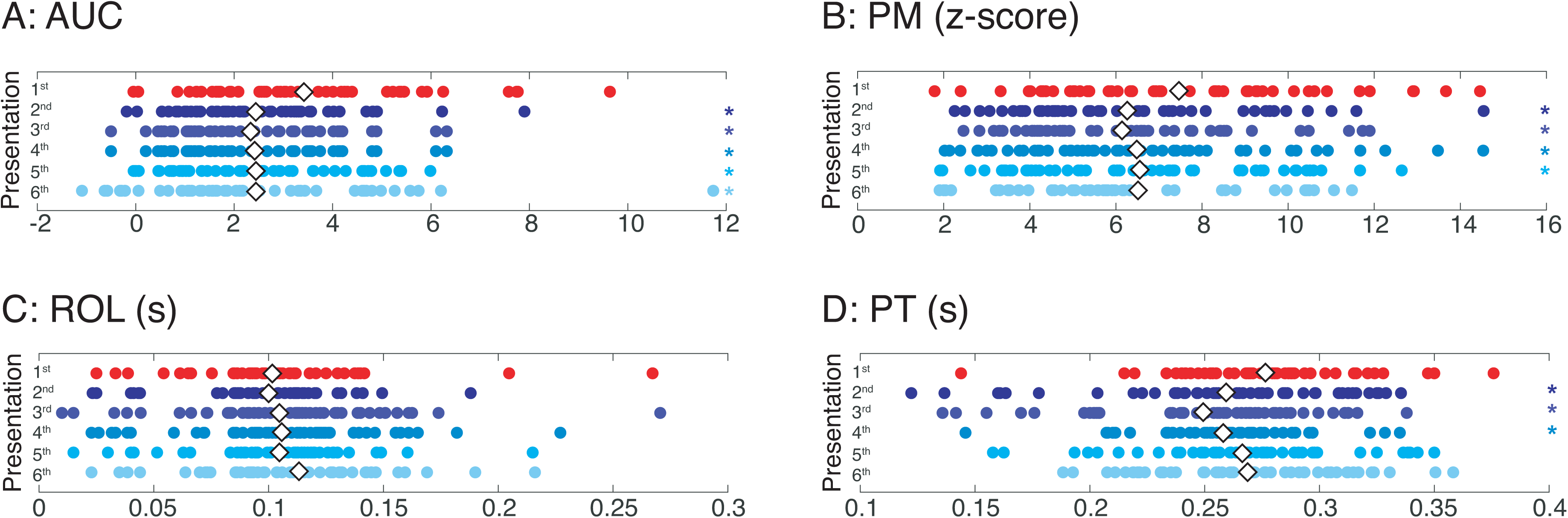
The effects of repetition on the magnitude and timing across multiple repetitions. The mean (A) AUC, (B) PM, (C) ROL, and (D) PT per electrode shown for the 1^st^ through 6^th^ repetition of faces. *Colored asterisk:* a significant difference (p<0.05, t-test) between that presentation number and the 1^st^ presentation. *White diamond*: mean value for each presentation number.

The mean AUC for subsequent face presentations was also smaller than the total response to the 1^st^ presentation of faces (1^st^= 3.4 ± 2.1; 2^nd^ = 2.5 ± 1.6; 3^rd^= 2.3 ± 1.5; 4^th^ = 2.4 ± 2.0; 5^th^ = 2.4 ± 1.7; 6^th^ = 2.4 ± 2.3). Statistical comparisons of the 1^st^ presentation versus 2^nd^ through 6^th^ presentations showed a significant reduction in the AUC of HFB responses (Figure 7A) with the reduction in the total response slightly increasing or remaining equal for presentations 2-6 compared to the 1^st^ presentation (t-test, FDR corrected: 1^st^ versus 2^nd^: t(47)= 4.16, p=1.34×10^−4^; 1^st^ versus 3^rd^: t(47)= 4.87, p=1.29×10^−5^; 1^st^ versus 4^th^: t(47)= 4.38, p=6.95×10^−5^; 1^st^ versus 5^th^: t(47)= 5.01, p=8.21×10^−6^; 1^st^ versus 6^th^: t(47)= 3.85, p=3.55×10^−4^). Notably, the 3^rd^, 4^th^, and 5^th^, but not 6^th^, presentations all showed a larger reduction in AUC than the reduction in AUC from the 1^st^ to 2^nd^ presentation.

Similarly, the PM of the response for subsequent face presentations was also smaller than the PM to the 1^st^ presentation of faces (1^st^ = 7.5 ± 3.0; 2^nd^ = 6.3 ± 2.7; 3^rd^ = 6.2 ± 2.4; 4^th^ = 6.5 ± 3.0; 5^th.^ = 6.6 ± 2.8; 6^th^ = 6.5 ± 3.8). Statistical comparisons of the 1^st^ presentation versus 2^nd^-5^th^ showed a significant reduction in the PM of HFB responses (Figure 7B, paired t-test, FDR corrected: 1^st^ versus 2^nd^: t(47)= 3.39, p=0.001; 1^st^ versus 3^rd^: t(47)= 3.67, p=6.25×10^−4^; 1^st^ versus 4^th^: t(47)= 3.06, p=0.004; 1^st^ versus 5^th^: t(47)= 2.98, p=0.005; 1^st^ versus 6^th^ Wilcoxon rank sum test: z=1.85, p=0.06). Notably, each of the 2^nd^, 3^rd^, 4^th^, and 5^th^ presentations generated a statistically significant reduction in PM, while the 6^th^ presentation did not generate a significant PM change compared to the 1^st^ presentation.

We next explored the temporal features of subsequent face repetitions. There was no statistical difference between the ROL for 1^st^ versus 2^nd^ to 6^th^ presentations (Figure 7C, paired t-test, FDR corrected: 1^st^ versus 2^nd^: t(47)=0.28, p=0.78; 1^st^ versus 3^rd^: t(47)= 0.27, p=0.78; 1^st^ versus sum test: z= −1.27, p=0.20). However, the mean PT for subsequent face presentations was generally earlier than the PT to the first presentation of faces (1^st^ = 275.9 ± 39; 2^nd^ = 259.1 ± 52; 3^rd^ = 249.0 ± 47; 4^th^ = 257.9 ± 33; 5^th^ = 265.9 ± 44; 6^th^ = 267.9 ± 42 ms). Statistical comparisons of the 1^st^ presentation versus the 2^nd^-6^th^ presentations, revealed an earlier PT of the HFB responses (Figure 7D) for the 2^nd^, 3^rd^, and 4^th^ presentations (paired t-test, FDR corrected: 1^st^ versus 2^nd^: t(47)=2.83, p=0.006; 1^st^ versus 3^rd^: t(47)= 4.43, p=5.63×10^−5^; 1^st^ versus 4^th^: t(47)= 3.38, p=0.001; 1^st^ versus 5^th^: t(47), p=0.10; 1^st^ versus 6^th^: t(47)=1.23, p=0.22). Notably, the 2^nd^, 3^rd^, 4^th^, presentations had a significantly earlier PT than the 1^st^ face presentation, with the maximum change for the 3^rd^ presentation.

Taken together, analyses of the dynamics of RS show that HFB responses across face repetitions are overall lower, have a smaller peak magnitude, and an earlier peak timing across multiple repetitions. These effects are summarized in **Table 2**. Our data suggest that RS effects depend on the number of repetitions, persist for the 3^rd^ and 4^th^ face repetitions, as well as show less consistent patterns for further repetitions. This indicates that repetition number and timing may be critical to the strength and nature of RS effects for long-lagged repetitions of faces in human VTC.

## Discussion

Using intracranial recordings, we characterized the effect of stimulus repetition on face-selective neural responses in human VTC for long-lagged stimulus repetitions (an average of 20.1 – 31.7s between repetitions) of faces and non-preferred categories. We found that RS occurs for high-frequency broadband (HFB) activity, but not lower frequency bands. Furthermore, we found that repetition reduces the total neural response (AUC) and peak magnitude (PM) in which more robust reductions occur in the former compared to the latter. RS effects occur early (within ~150 ms after stimulus onset) and persist for the entire stimulus duration. Interestingly, HFB responses had different temporal characteristics for subsequent than initial face presentations: neural responses were faster to peak for the 2^nd^ versus 1^st^ presentation, even though there was no significant differences in response onset timing. This response characteristic did not occur for other non-preferred categories, but consistently occurred for additional face repetitions with additional intervening stimuli. In the sections below, we further discuss RS effects across frequency bands, as well as theoretical models of RS.

### Repetition Suppression in HFB, but not Lower Frequency Bands

Our data reveal RS in HFB (>70Hz), but not lower frequency bands (<30Hz). These findings are consistent with ECoG data from Engell AD and G McCarthy (2014) who showed that for immediate repetitions of faces (8 repetitions of the same face, 2s ISI, and no intervening stimuli), there were decreases in responses in the low (30-60Hz) and high gamma range (60- 100Hz) with no significant changes in the α(8-12Hz) or β (15-30Hz) range. As lower frequency activity is associated with visual evoked potentials (VEP) (Allison T *et al.* 1999; Rossion B 2014), and VEP and HFB responses are often dissociated in face-selective regions (Engell AD and G McCarthy 2011; Rangarajan V *et al.* 2014), our observations are in line with prior studies that measured VEPs and failed to find RS to faces in ECoG data from VTC (Puce A *et al.* 1999). Specifically, the lack of RS in lower frequencies in the present study and the coupling between VEP and low frequency activities is consistent with the lack of RS in the intracranial VEP observed in prior studies. It is possible that additional effects of repetition may be present in the scalp VEP (Schweinberger SR *et al.* 1995; Doniger GM *et al.* 2001; Schweinberger SR *et al.* 2002; Kuehl LK *et al.* 2013) or in synchronous or phase-locked features of low frequency responses (Gilbert JR et al. 2010; Engell AD and G McCarthy 2014), but we have not observed such effects in the current study. Nonetheless, we find robust RS to faces early in the HFB response and in the majority of face-selective electrodes indicating that future studies of stimulus repetition using intracranial measurements should quantify HFB responses.

Our finding of RS in HFB activity is also consistent with prior studies showing RS in gamma and high-gamma bands of local field potentials (LFP) for repetitions of faces (Engell AD and G McCarthy 2014), letters (Rodriguez Merzagora A *et al.*), and objects (McMahon DB and CR Olson 2007; De Baene W and R Vogels 2010; Friese U et al. 2012; Engell AD and G McCarthy 2014) in both human VTC as well as macaque inferior temporal cortex. Additionally, we found that significant RS occurs within 350 ms of stimulus onset and persists for the remainder of the image duration. As HFB activity is correlated with local neuronal firing (Logothetis NK *et al.* 2001; Nir Y et al. 2007; Miller KJ *et al.* 2010), our results suggest that RS in HFB is due to reduced neuronal firing rate to repeated stimuli, which is consistent with findings of RS in single neurons of macaque inferior temporal cortex (Miller EK *et al.* 1991; Li L et al. 1993; McMahon DB and CR Olson 2007; De Baene W and R Vogels 2010).

### Implications of our findings on theoretical models of repetition suppression

Several theories have been proposed to account for RS in high-level visual cortex such as scaling, sharpening, facilitation, synchrony, prediction error, and combinations therein (Grill-Spector K *et al.* 2006; Summerfield C et al. 2008; De Baene W and R Vogels 2010; Weiner KS *et al.* 2010; Gotts SJ et al. 2012; Henson RN 2016; Vogels R 2016; Alink A *et al.* 2018). To date, most empirical data testing these theories have largely examined the effect of repetition on the magnitude of responses, as the prevalent scaling and sharpening models make specific predictions about the amplitude, rather than timing of RS.

Both scaling and facilitation models of RS predict the largest reduction in amplitude for repetitions of the preferred stimulus, while sharpening predicts the smallest reduction in amplitude to repetitions of the preferred stimulus. Consistent with scaling and facilitation models, we find lower PM and AUC for the 2^nd^ versus the 1^st^ presentation of a face. Notably, we also find the largest RS for the preferred stimulus (faces) as compared to non-preferred stimulus categories, consistent with prior fMRI studies (Weiner KS *et al.* 2010). As repetitions of the same stimulus occurred with long lags and many intervening stimuli, these RS effects are impressive, and argue against the hypothesis that reduced responses to repeating stimuli stem from low-level, image-based adaptation effects.

The facilitation model is the only model of RS that makes predictions about the temporal characteristics of repetition effects. At its simplest, facilitation predicts that repetition causes faster processing of stimuli, that is, shorter latencies or shorter durations of neural firing (Grill-Spector K *et al.* 2006). However, researchers vary in their predictions of the temporal dynamics associated with facilitation models ranging from: (a) a faster response onset latency (ROL) (James TW and I Gauthier 2006), (b) no change in the ROL (Henson RN 2012, 2016), (c) an earlier peak time (James TW and I Gauthier 2006), or (d) a shorter response duration (Henson RN and MD Rugg 2003; Henson RN 2016) for the repeated presentation of an image. Intriguingly, our results provide the first empirical support for the timing predictions of the accumulation model of James TW and I Gauthier (2006), which is an extension of the facilitation model, during long-lagged stimulus repetitions as the 2^nd^, 3^rd^, and 4^th^ repetition of faces produced neural responses that were faster to peak than the 1^st^ presentation. We find that in face-selective electrodes, temporal changes in the HFB responses to repeated stimuli were unique to faces. This suggests that in a stimulus-selective cortical region, temporal facilitation in peak timing may occur for the preferred stimulus, but not to non-preferred stimuli. As recent experiments have shown that modeling the temporal responses of neurons in millisecond resolution better predicts BOLD responses (with a resolution of seconds) than the general linear model (Stigliani A et al. 2017; Zhou J et al. 2018; Stigliani A et al. 2019; Zhou J et al. 2019), future research can measure both ECoG and fMRI responses in an event-related design for long-lagged repetitions to directly relate the impact of the repetition on the combined changes to PT, PM, and AUC, measured with ECoG, on BOLD responses.

Other studies have suggested that RS is linked to predictability (Summerfield C *et al.* 2008). We believe that predictability is an unlikely explanation underlying the different temporal features of neural responses during repeated compared to non-repeated conditions in the present study for five reasons. First, our long-lagged experimental design controlled for stimulus predictability: there was an equal occurrence of repeated and non-repeated images throughout the experiment, and participants were unaware when repetitions were going to occur. Second, subjects participated in an orthogonal fixation task, which did not require judgment of faces or top-down attention to faces. Third, while RS is a reliable effect across studies, stimuli, and measurements (Grill-Spector K *et al.* 2006; Summerfield C et al. 2008; De Baene W and R Vogels 2010; Weiner KS *et al.* 2010; Gotts SJ et al. 2012; Henson RN 2016; Vogels R 2016; Alink A *et al.* 2018), the effects of stimulus predictability on the magnitude of RS are inconsistent across studies, stimuli, and measurements (Sawamura H et al. 2006; Kaliukhovich DA and R Vogels 2011; Kovacs G et al. 2013; Tang MF et al. 2018; Vinken K et al. 2018). Fourth, RS to repetitions of non-preferred stimuli did not always occur within face-selective electrodes even though the repetition predictability of non-preferred stimuli was equally as likely as face repetitions. Fifth, to our knowledge, the prediction error account does not make explicit predictions regarding the temporal dynamics of responses, such as the faster time to peak for repetitions of preferred stimuli that eventually plateaus after the 4^th^ repetition. Thus, while the prediction error account for RS is an appealing hypothesis, it does not explain RS during long- lagged paradigms with intervening stimuli.

### Theoretical and Methodological Implications for Future Work

A theoretical implication resulting from comparing our findings to previous work is that different experimental paradigms may affect different aspects of the temporal dynamics of neural responses in VTC. For example, while we found no effect of repetition for ROL, previous intracranial studies recording from VTC in humans (Engell AD and G McCarthy 2014; Rodriguez Merzagora A *et al.* 2014) and non-human primates (McMahon DB and CR Olson 2007; Anderson B et al. 2008; De Baene W and R Vogels 2010; Engell AD and G McCarthy 2014) have shown delayed ROL. As such, future modeling work should take into consideration how the nature of stimulus repetition may affect the temporal properties of the response such as ROL and PT. For instance, previous work has reported delayed ROL in VTC for immediate stimulus repetitions with no intervening stimuli and less than 2 seconds between repetitions implemented in short-lagged paradigms (Anderson B *et al.* 2008; De Baene W and R Vogels 2010; Engell AD and G McCarthy 2014; Rodriguez Merzagora A *et al.* 2014). This suggests that immediate repetitions and long-lagged repetitions with intervening stimuli (as in the present study) may have different temporal characteristics. For example, using letters during a Sternberg working memory task and ECoG in human occipital and temporal cortex, Rodriguez Merzagora A *et al.* (2014) and colleagues reported RS only for HFB, but unlike the present study, found that the ROL for repeated stimuli was actually slower than for non-repeated stimuli. The combination of the present and past findings further suggests the possibility that different neural mechanisms may lead to the differential temporal effects across repetitions given varied stimulus timing parameters. This idea is particularly interesting given that long-lagged repetitions reflect an implicit memory trace while immediate repetitions may be related to refractory periods that may dampen the generation of action potentials (Grill-Spector K *et al.* 2006; Fabbrini F *et al.* 2019). Future studies can test this hypothesis by systematically varying the interval between repetitions of the same stimuli, measuring how timing properties affect the temporal characteristics of the neural response, and further applying computational encoding models to understand the temporal features of stimulus repetitions (Stigliani A *et al.* 2019). Additionally, as RS effects vary across brain regions (Verhoef BE et al. 2008; Weiner KS et al. 2010), future studies are needed to a) determine if the facilitation of the PT observed here is prevalent across other regions or is specific to human VTC, b) explore the dynamics of repetition effects using other visual categories and for non-visual domains, and c) examine how the number of intervening stimuli and interstimulus interval may affect RS. Such work could also explore whether information can be classified more quickly for repeated versus novel presentations of stimuli across visual cortex intracranially (Weiner KS *et al.* 2010).

### Open Questions for Future Research

Our data provide the first insights of the effect of long-lagged repetitions on HFB responses in VTC. However, a few questions are unanswered by the present study and can be addressed in future studies.

First, why wasn’t RS to long-lagged stimulus repetitions observed in all electrodes? Our data revealed that 3 electrodes showed a weak enhanced response to the 2^nd^ versus 1^st^ presentation. We note that the timing of enhancement in these electrodes is very different than those showing RS: the first time point showing significant repetition enhancement appeared later in the trial (on average ~375 ms) as compared to significant repetition suppression (on average ~150 ms) after stimulus onset. These varied temporal profiles of repetition suppression versus repetition enhancement may suggest that different mechanisms underlie these phenomena. For example, bottom-up signals may drive repetition suppression whereas top-down feedback signals may drive repetition enhancement (Cheal M et al. 1991; Pinto Y et al. 2013). These hypotheses can be examined in future work.

Second, why was the shift in peak timing for face repetitions not observed in all electrodes? This may be in part due to the varied anatomical location of our face-selective electrodes or the initial strength of the face responses in these electrodes. Future studies with more face-selective electrodes are necessary to investigate channels that show a varied temporal profile. This would also help address inter-subject variability inherent to intracranial research (Aarts E et al. 2014). A larger pool of subjects, with more face-selective electrodes, would allow us to further address how individual differences may influence RS effects.

Third, how does the neural response magnitude to preferred stimuli affect RS? Previous electrophysiology studies in macaque IT (Sawamura H *et al.* (2006) have explored the relationship between response strength and repetition. Here, we show that the effects of repetition are larger for the preferred category (faces) than non-preferred categories (cars, houses, limbs) consistent with fMRI data and with the scaling model of RS (Weiner KS *et al.* 2010). One appealing approach to address this question in future research may be to use presentation methods that modulate responses to faces such as lowering their contrast or embedding them in noise to examine the effect of level of response on the timing of neural responses to repeated stimuli.

### Conclusion

In sum, neural RS is a pervasive effect across high-level visual regions in cortical sensory systems. Understanding and characterizing RS dynamics – across temporal and neuroanatomical spatial scales – serves as a window to understand how cortical responses depend on the history of a stimulus, which has direct implications for modeling neural systems subserving learning, memory, and perception. Here, we leveraged the precise spatial localization and high temporal resolution of ECoG from 9 human subjects implanted with intracranial electrodes in VTC to measure the effects of stimulus repetition after tens of seconds. Our approach revealed that RS (a) occurs in VTC activity in HFB range, but not lower frequency bands, (b) is associated with lower peak magnitude, lower total responses of the HFB signal, and earlier peak responses, and (c) RS effects occur early (~150 ms) and persist for the entire stimulus duration. Furthermore, these temporal dynamics of RS were largely specific to the preferred stimulus (faces) within face-selective electrodes in VTC and persisted across additional repetitions until about the 4^th^ repetition. Together, these data motivate future empirical and theoretical work examining the effects of stimulus repetition on neural responses not only within VTC, but also throughout the brain and in other sensory and non-sensory domains.

## Acknowledgements

All intracranial data were acquired at Stanford Medical Center through collaboration with Professor Josef Parvizi, MD PhD at the Laboratory of Behavioral and Cognitive Neuroscience (LBCN). We thank the patients for participating in our research; Dr. Brett L. Foster, Dr. Jesse Gomez, and Sandra Gattas for help with data collection; and Dr. Nathan Witthoft for coding the experiment.

## Support

National Science Foundation Graduate Research Fellowship to V.R. (DGE 1106400); National Institute of Neurological Disorders and Strokes at the National Institutes of Health (grant number R37NS21135) & The James S. McDonnell Foundation Grant to RTK; National Eye Institute at the National Institutes of Health (grant number 1R01EY02391501A1) to KGS

## Author Contributions

CJ, KSW, KGS designed the experiment; VR, CJ, KSW collected the data; VR, RTK, KSW, KGS contributed to data analysis; VR, KSW, KGS wrote the manuscript; All authors approved the manuscript.

---

**Movie 1: The temporal dynamics of average RS in each of the 48 face-selective electrodes are presented on the MNI average brain.**

The spatial-temporal distribution of RS is visualized and corrected for electrode overlap density. For display purposes, the additive spatial distribution of *t*-values are scaled to the same maximum (2, red) and minimum (−2, blue), as well as corrected for the spatial overlap of included electrodes. Electrode locations for all subjects are overlaid on the MNI brain to display the temporal evolution of RS. The video is slowed down 10 times for ease of viewing.

**Table 1:** Patient information including subject (S1-S9), electrode number, MNI coordinates (X,Y,Z mm) for each face-selective electrode, fixation task performance (hit rate and mean response time), and significance value of the AUC reduction between the 2^nd^ and 1^st^ presentation (t-test, FDR corrected across trials).

**Table 2:** The effects of repetition on both magnitude (AUC, PM) and timing (ROL, PT) are summarized for the 1^st^ presentation versus 2^nd^ – 6^th^ presentations. Bolded values indicate significant differences (p<0.05, t-test) between that presentation number and the 1^st^ presentation.

**Supplemental Figure 1:**
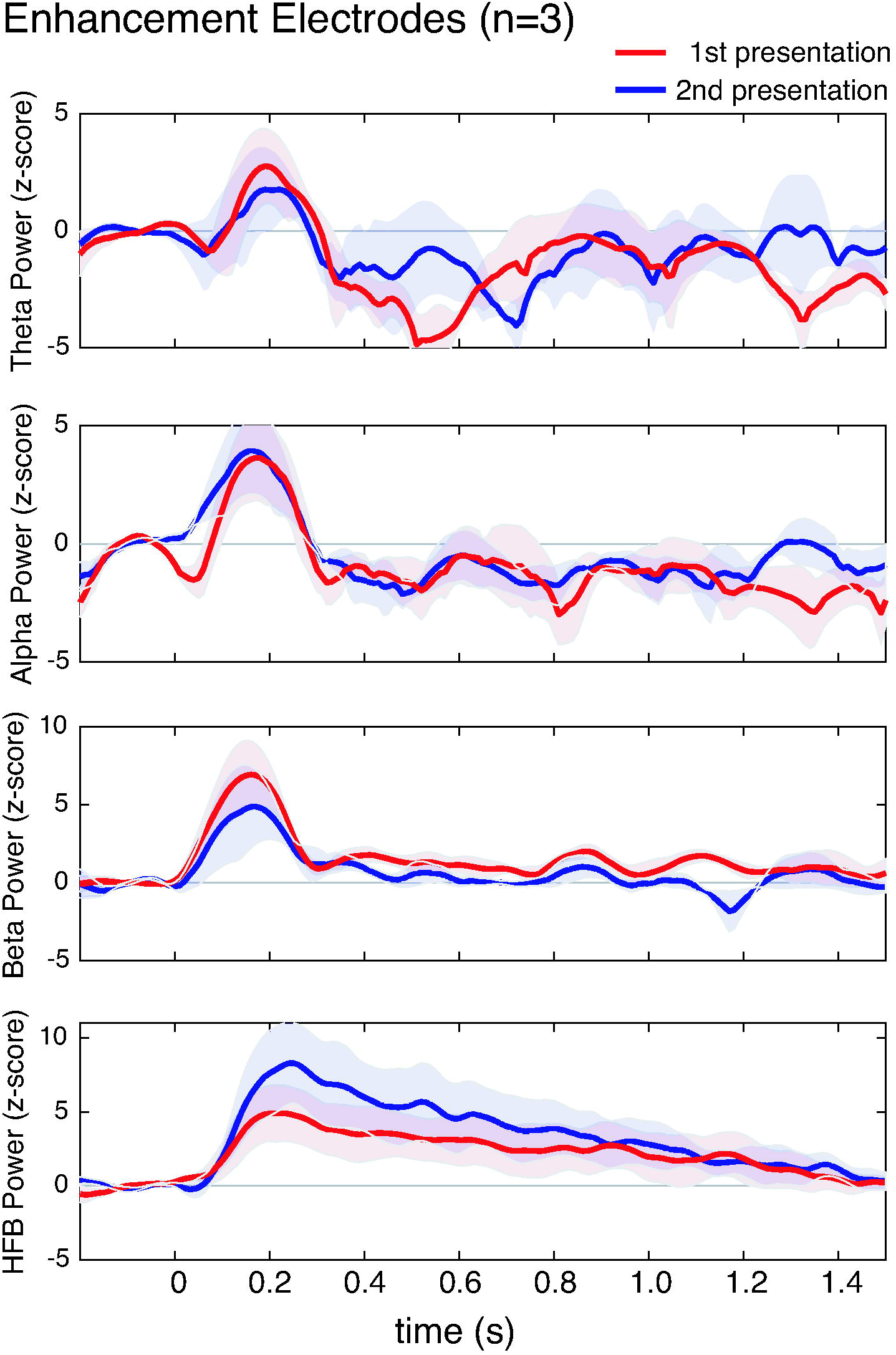
Repetition enhancement in human VTC. Average responses to 1^st^ (red) versus 2^nd^ (blue) presentations of faces across 3 face-selective electrodes that showed significant HFB repetition enhancement. Each row shows responses in a different frequency band. *From top to bottom*, frequencies are: theta (Θ: 4-8 Hz), alpha (α: 8-13 Hz), beta (β: 16-30 Hz), and high frequency broadband (HFB: 70-150 Hz). *Shaded region:* standard error of the mean across 3 face-selective electrodes.

**Supplemental Figure 2:**
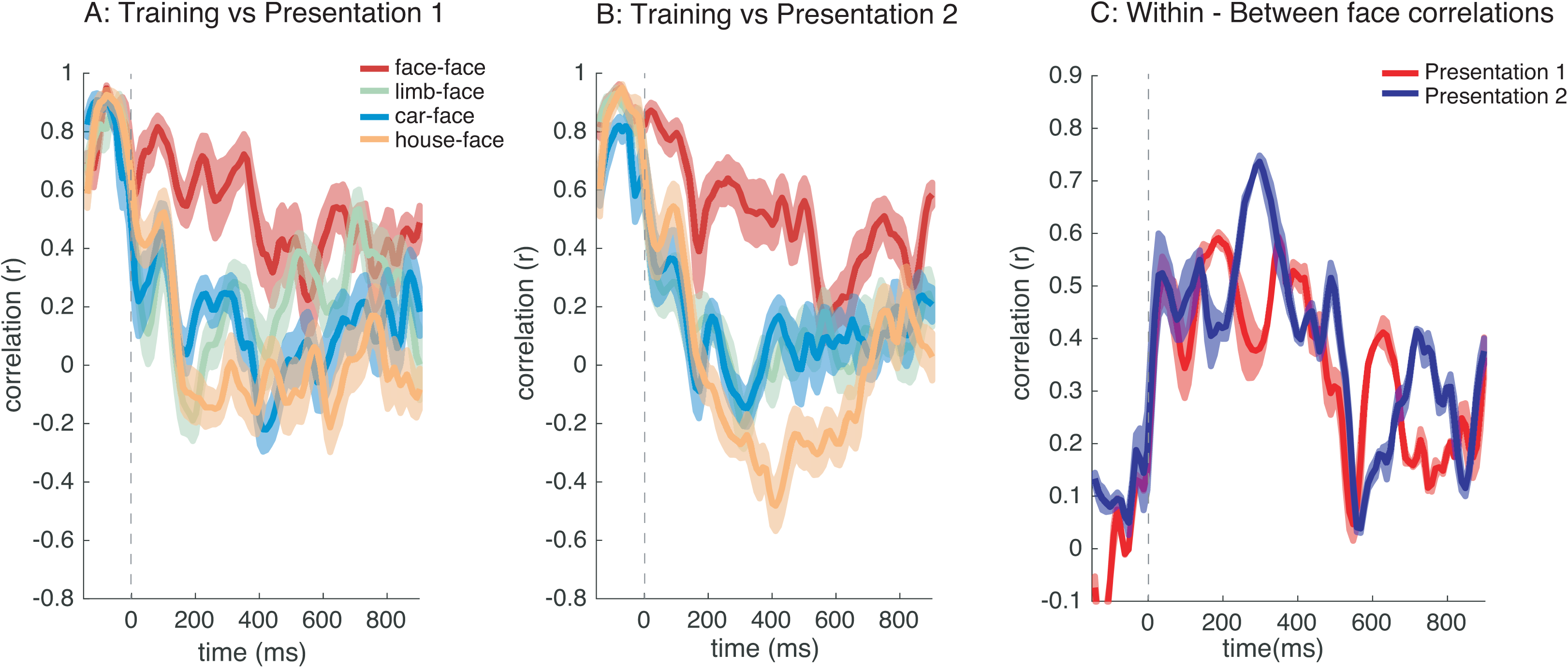
Analysis of distributed responses to the four categories (face, limb, car, house) over time and repetitions. The correlation between the distributed responses of training images (non-repeated faces) and each of the testing sets across all 48 face-selective electrodes as in Figure 6: (A) 1^st^ and (B) 2^nd^ face and non-preferred categories. Color indicates the categories that are correlated: *red:* face-face; *orange:* houses-face; *green:* limbs-face; *blue:* cars-face correlations, respectively). (C) Within-face correlation minus between mean face and non-face correlations for 1^st^ and 2^nd^ presentations are plotted over time; dashed vertical lines: stimulus onset*. Error bars: indicate standard error across bootstraps*

**Supplemental Figure 3:**
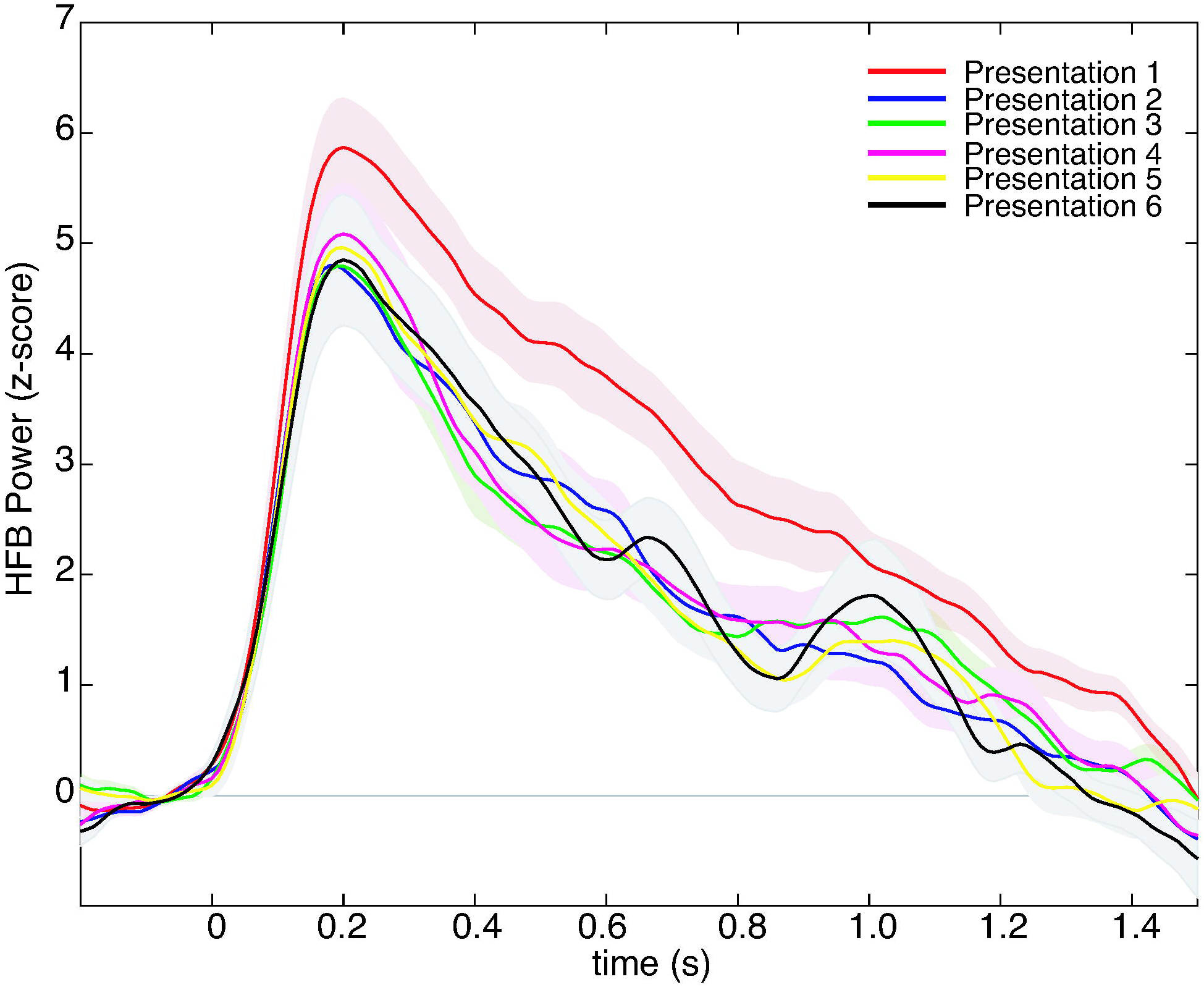
Average HFB timecourse of 6 face repetitions across all face-selective electrodes. Mean HFB responses to faces across 48 face-selective electrodes for each repetition number. Color indicates repetition number. Shaded regions indicate the standard error of the mean.

## Notes

### Competing Interest Statement

The authors have declared no competing interest.

### Summary of Updates

This article has been accepted for publication in Cerebral Cortex published by Oxford University Press.

